# Deep coverage microscopy exposes a pharmacological window for modifiers of neuronal network connectivity

**DOI:** 10.1101/555714

**Authors:** Marlies Verschuuren, Peter Verstraelen, Gerardo Garcia, Ines Cilissen, Emma Coninx, Mieke Verslegers, Peter Larsen, Rony Nuydens, Winnok H. De Vos

**Affiliations:** Laboratory of Cell Biology & Histology, University of Antwerp, Department of Veterinary Sciences, Antwerp, Belgium; Animal Physiology and Neurobiology, KU Leuven, Department of Biology, Antwerp, Belgium; Radiobiology Unit, Institute of Environment, Health and Safety, Belgian Nuclear Research Centre, Mol, Belgium; Janssen Research & Development, a division of Janssen Pharmaceutica NV, Beerse, Belgium; University of Ghent, Department of Molecular Biotechnology, Ghent, Belgium

**Keywords:** synaptic connectivity, neuronal network morphology, calcium imaging, high-content screening, neurodegeneration, antioxidant depletion, hTau.P301L

## Abstract

**Background:** Therapeutic developments for neurodegenerative disorders are redirecting their focus to the mechanisms that contribute to synaptic plasticity and the loss thereof. Identification of novel regulators requires a method to quantify neuronal network connectivity with high accuracy and throughput. To meet this demand, we have established a microscopy-based pipeline that integrates morphological and functional correlates of connectivity in primary neuronal culture.

**Results:** We unveiled a connectivity signature that was specific to the cell type and culture age. We defined a score that accurately reports on the degree of neuronal connectivity and we validated this score by targeted perturbation of microtubule stability and selective depletion of anti-oxidants. With a focused compound screen, we discovered that inhibition of dual leucine zipper kinase activity increased neuronal connectivity in otherwise unperturbed cultures and exerted neuroprotective effects in cultures grown under sub-optimal or challenged conditions.

**Conclusions:** Our results illustrate that profiling microscopy images with deep coverage enables sensitive interrogation of neuronal connectivity and allows exposing a dose and time window for pharmacological interventions. Therefore, the current approach holds promise for identifying pathways and compounds that preserve or rescue neuronal connectivity in neurodegenerative disorders.

## Background

Wiring the central nervous system demands precise formation and maintenance of neuronal connections. Functional connections between neurons are established by means of micron-sized interfaces, called synapses [1,2]. Synaptic activity and the adjoined opening of gated calcium channels generate calcium transients that drive well-known morphological changes such as dendritic growth and arborization, but which can also influence synapse strength [3]. This dynamic remodeling of both neurites and synapses fosters improved communication between neurons allowing synchronous functional activity, thereby reinforcing the overall connectivity of the neuronal network. All long-lasting adaptations of the brain, including learning, memory, addiction and chronic pain sensation, rely on the continuous remodeling of neuronal network connectivity [3]. And, disruption of this process is a hallmark of numerous neurological diseases, including schizophrenia, major depressive disorder and Alzheimer’s disease (AD) [4]. For example, the cognitive impairments witnessed in AD patients correlate with synapse and dendritic loss as well as a reduction of the brain activity, indicating an overall decrease in neuronal connectivity [5-7]. Therefore, therapeutic developments for neurodegenerative disorders focus on identifying regulators that promote neuronal connectivity or prevent the loss thereof.

The dense, three-dimensional organization and spatial heterogeneity of the brain makes studying neuronal connectivity *in vivo* a daunting task, which is not amenable to upscaling. Therefore, systematic screening efforts most often rely on simplified models such as neuronal cell cultures. Although some immortalized tumor cells (*e.g.*, SH-SY5Y human neuroblastoma cells, NT2 human teratocarcinoma cells, PC12 rat pheochromocytoma cells of the adrenal medulla) can be differentiated to take on a neuron-like phenotype (*e.g.*, neurite outgrowth, expression of synaptic markers, induced or subtle spontaneous calcium activity), none fully recapitulate the full feature set of a physiologically connected neuronal network [8-13]. Neurons derived from human induced pluripotent stem cell have a much higher translational value in comparison with other cell lines and primary cultures, but the differentiation process is very labor and time expensive [14-16]. That is why, as yet, primary neuronal cultures represent the model of choice for genomic and pharmacological high-throughput screens [17-25]. However, most of these screens tend to focus on one or two specific readouts such as neuron number [19,24], neurite outgrowth [18,19,24,25] or synapse density [22,23]. This reductionist approach can be misleading since it neglects multifactorial effects. For instance, changes in synapse density can depend on alterations in the actual number of synapses [20,21], but could also be influenced by changes in the neurite density, without affecting the actual synapse count [26]. Furthermore, solely gauging morphological correlates may mask potential changes in functional connectivity. Indeed, it has already been shown that spontaneous activity in primary cultures does not scale linearly, but rather exponentially, with synaptic density [27], and that morphological aberrations of primary neuronal networks do not always result in functional impairments [26]. In order to accurately quantify the overall neuronal connectivity in primary cultures, information gained from several readouts should ideally be combined. Hence, instead of using one or a few descriptors, here we comprehensively assess the major morphological and functional correlates of neuronal network connectivity, and we integrate them to accurately map changes between subsequent maturation stages *in vitro*. In a targeted compound screen, we identified dual leucine zipper kinase (DLK) inhibition as a positive modulator of neuronal connectivity in unperturbed cultures and as a neuroprotector in cultures grown under suboptimal or compromised conditions.

## Results

### Culture age correlates with both morphological and functional changes in primary neuronal networks

When cultured *in vitro*, primary neurons form dense dendritic networks and develop functional activity, epitomized by a synchronicity of intracellular calcium fluctuations [27-29]. To determine the temporal evolution of these changes, we first cultured cortical neurons for 48 days *in vitro* (DIV), and quantified a variety of morphological parameters at 6-day intervals (Fig. 1a). We thereby made a distinction between descriptors that inform on one of three major categories, namely the dendrite network, the synapse markers, or the nuclei (Additional file 1: Table S1). Up to 36 DIV, the neuronal network became denser, as exemplified by an increase in dendrite density (the area of the field of view covered by dendrites) and the number of nodes (branch points) in the network (Fig. 1b). At later time points, the dendrite network deteriorated. The presynaptic spot density – inferred from Synapthophysin immunolabeling – increased rapidly up to 18 DIV, stabilized up to 30 DIV and eventually decreased. The postsynaptic spot density – as gauged from PSD-95 immunolabeling – followed a more gradual evolution and reached a maximum at 42 DIV. In line with this, the relative number of overlapping pre- and postsynaptic spots, a proxy for synapse density, stabilized at 18 DIV (Fig. 1b). The number of neuronal nuclei (and thus neurons) gradually decreased with culture age. The ratio of neuronal cells versus non-neuronal (glial) cells remained relatively constant until 36 DIV, after which it significantly decreased, predominantly due to a stronger loss of neurons *vs.* glial cells (Fig. 1b). Thus, this multiparametric analysis showed that cortical neurons develop progressive morphological connectivity up to 36 DIV with the strongest evolution taking place between 3 and 18 DIV.

**Fig 1.**
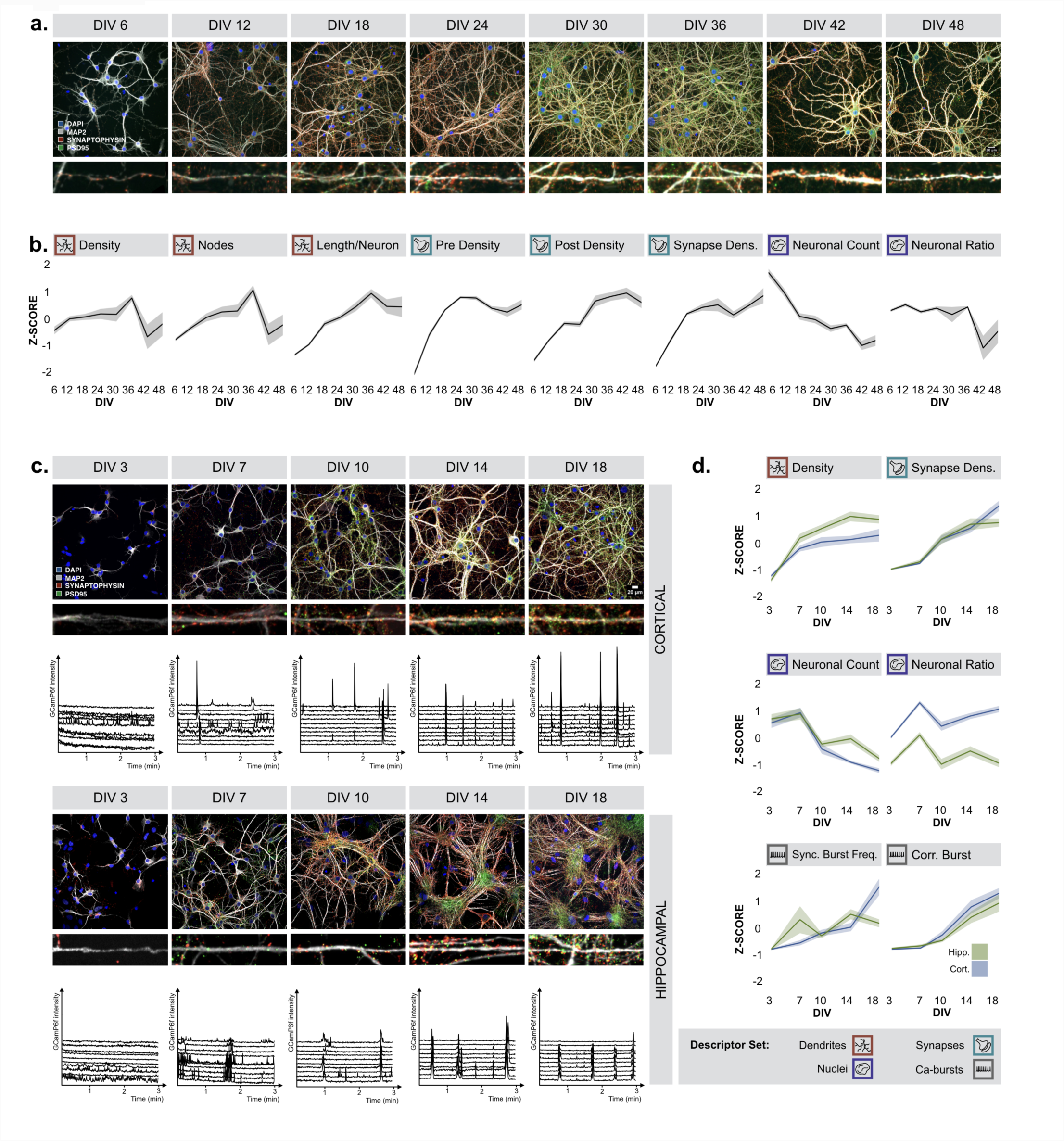
Culture age correlates with morpho-functional changes. **(a)** Cortical cultures were grown for an extended time period to quantify morphofunctional changes. Representative images of cortical cultures fixed at 6, 12, 18, 24, 30, 36, 42 and 48 DIV are visualised. **(b)** Quantification results show that the dendrite density and nodes increased gradually untill 36 DIV, after which they decreased. The deterioration of the dendrite network was less pronounced when normalized to the number of neurons which decreased over time. The presynaptic density and synapse density increased during the first 18 DIV. The density of postsynaptic spots increased gradually. The ratio between neuronal and nonneuronal nuclei remained more or less the same until 36 DIV (n_bio_=2 x n_tech_=6). (**c**) Representative images and Ca^2+^- traces (10 neurons) for a more resolved follow-up experiment (3,7,10,14,18 DIV) of primary hippocampal and cortical cultures. **(d)** Similar trends are observed in selected morphological descriptors (n_bio_=3 x n_tech_ =6). Synchronous activity increased from DIV 10 onwards (n_bio_=3 x n_tech_ =6).

We next wondered whether this evolution was generic to cultures derived from different brain regions. Therefore, we compared cortical with hippocampal cultures and quantified changes with a higher time resolution (3, 7, 10, 14 and 18 DIV) (Fig. 1c). For both neuronal culture types, we found similar trends in the previously described descriptors (*e.g.*, dendrite density, synapse density, nuclear count) (Fig. 1d), suggesting that they both become morphologically more connected during this culture period. However, there were also culture type-dependent differences. Hippocampal neurons displayed greater tendency to form tight dendrite bundles than cortical neurons. This clustering occurred especially at later time points (Fig. 1c and Additional file 2: Figure S1). Hippocampal cultures also showed a stronger postsynaptic marker intensity, whereas nuclei from cortical neurons displayed more pronounced chromocenters contributing to a stronger spot-like texture. Arguably one of the most striking differences between both cultures, was the ratio of neuronal cells versus non-neuronal cells, which was far higher in cortical cultures, due to a comparatively low number of glial cells (with larger, flattened nuclei, resulting in a lower average nuclear area) (Additional file 2: Figure S1).

Recognizing that morphological measures do not necessarily report on functional changes in connectivity [26], we complemented the morphological analysis with a previously optimized live cell calcium imaging assay [30]. A very similar evolution was found for both culture types, with virtually no synchronous bursting activity - expressed as the correlation between calcium bursts of all active neurons in a field of view - at 3 or 7 DIV, and progressively more synchronous activity at later time points (Fig 1d). Thus, we conclude that primary hippocampal and cortical cultures form neuronal networks that become both morphologically and functionally more connected with culture age, at least up to 18 DIV.

### Neuronal culture states can be distinguished by their morphofunctional signature

Having established that both hippocampal and cortical neurons showed distinct, yet consistent morphofunctional changes with culture age, we asked whether we could retrieve this evolution in an unsupervised manner from the extracted descriptors. Hierarchical clustering on the entire data set revealed that each condition had a distinct descriptor profile, whereby many descriptors showed a high correlation with culture age (47 % of the descriptors had an absolute correlation with DIV > 0.5) (Fig. 2a). We then applied principal component analysis (PCA) to the morphological descriptor set. Despite significant variability between individual biological replicates (Additional file 3: Figure S2a), the combined data showed a clear pattern in PC space: when plotting the two first principal components, each cell type displayed a distinct sequence of clusters following a trajectory according to the culture age (Fig. 2b). We also tested PCA on the separate morphological descriptor classes (nuclei, neurite, synapse), but no single class could unequivocally discriminate both cell type and culture time as good, indicating that a combined descriptor set yielded the most powerful discriminatory fingerprint (Additional file 3: Figure S2b). PCA of the functional descriptor set did not lead to well-defined trajectories either, most likely due to the fact that synchronous activity only surfaced from DIV 10 onwards (Fig 2b). Given the clear separation in descriptor space based on PCA, we next asked whether we could use this information to predict culture age per cell type. To this end, we trained a random forest classifier (RFC) using different parameter settings and selected the best performing classifier using 10-fold cross-validation on a training set, after which we determined the misclassification rate (MCR) on a test set (Fig. 2c). A classifier based on all morphological descriptors yielded a MCR of 5%, while the individual descriptor sets yielded classifiers with MCRs above 14% (Additional file 3: Figure S2c). Not unexpectedly, the performance of a classifier that was solely based on functional descriptors was poor (MCR > 45%). Yet, the fact that a morphological descriptor set could be used to untangle and even predict culture age, suggested that an integrated signature could be indicative of the degree of neuronal connectivity.

**Fig 2.**
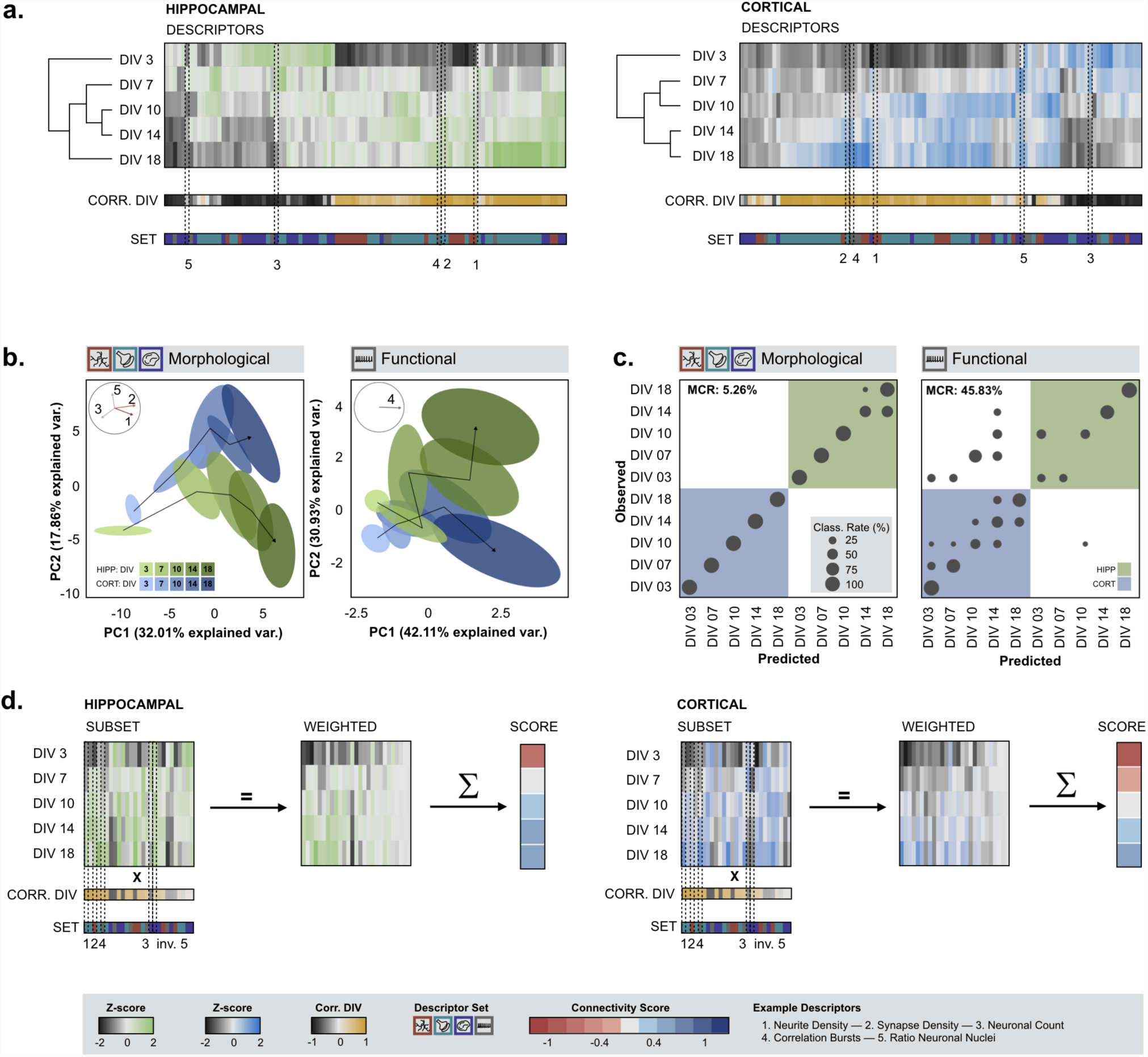
Cultures can be clustered and classified based on morphofunctional signature. **(a)** Hierarchical clustering based on z-scores of all descriptors. The correlation of the descriptors with culture age is indicated (black/orange color coded) as well as the descriptor set to which they belong. Example descriptors, discussed in Fig. 1 are indicated. **(b)** PCA based on morphological descriptors distinguished hippocampal from cortical cultures. Data points clustered according to culture time and followed distinct temporal trajectories along the direction of a decreasing nuclear count and increasing dendrite- and synapse density (n_bio_=3 x n_tech_ =6). Trajectories formed by the PCA of functional descriptors could only differentiate early (3-10 DIV) from later time points (14-18 DIV) (n_bio_=3 x n_tech_=6). (**c**) Confusion matrices and classification results using a random forest classifier. The misclassification rate (MCR) of the classification based on morphological descriptors was 5% and much lower than the MCR of the classification based on functional data (45%). (**d**) The z-scores of a subset of descriptors were multiplied with their respective correlation with culture age (weighted z-scores) and summed to obtain a connectivity score that allows quick interpretation of the connectivity state at a given timepoint.

Due to a different experimental setup (and number of technical replicates), morphological and functional data could only be combined at the level of the biological replicate. This reduced the number of data points drastically and precluded reliable PCA or RFC. However, we reasoned that the exclusion of functional descriptors from a connectivity analysis could lead to a bias under specific challenged conditions. Indeed, when we treated cortical cultures with the N-methyl-d-aspartate sensitive glutamate receptor (NMDA-R) antagonist MK-801, we found an adverse impact on functional connectivity that was significant at 18 DIV (p<0.05, pairwise Wilcoxon test with Bonferroni correction), without showing significant changes in key morphological descriptors (p>0.05, pairwise Wilcoxon test with Bonferroni correction) (Additional file 4: Figure S3). The same holds true for nuclear descriptors, which at first sight may seem less relevant for assessing culture age. Yet, a chronic incubation with arabinosylcytosine (AraC, which selectively blocks astrocyte proliferation) had a significantly negative impact on the non-neuronal cell number during the whole time range, (p<0.05, pairwise Wilcoxon test with Bonferroni correction), without affecting the synapse density (p>0.05, pairwise Wilcoxon test with Bonferroni correction) (Additional file 5: Figure S4). Therefore, we sought an approach to integrate all morphofunctional descriptor classes in such a way that it intuitively reports on connectivity changes without disregarding putative off-target effects (Fig. 2d). To avoid redundancy, we excluded descriptors that showed high inter-correlation (> 0.75), whilst giving priority to descriptors that showed the highest correlation with the culture age (Additional file 6: Figure S5). From this subset, a weighted average was calculated using the correlation with culture age (DIV) as weights. This resulted in a single metric which we refer to as connectivity score (Fig. 2d). The connectivity score is sensitive to changes in dendrite, synapse and nuclear descriptors, as well as functional descriptors, which makes the connectivity score a robust metric that captures the degree of connectivity in a specific condition (Additional file 7: Figure S6).

### Focused compound screen identifies DLK as a positive modulator of neuronal network connectivity

Using the connectivity score as readout, we subsequently initiated a compound screen to expose regulators of neuronal network connectivity (Fig. 3). Given the higher culture yield, better reproducibility, and lower tendency to cluster, we chose to continue with cortical cultures. A rational selection of putative molecular targets (mTOR, NMDA-R, histone deacetylases (HDAC), DLK) was made based on their positive effect in neurodegenerative conditions described in literature and for each target, at least one compound (rapamycin, memantine, MK-801, suberoylanilide hydroxamic acid (SAHA), tubastatin, GNE3511) was selected [31-37]. Rapamycin, inhibitor of mTOR, was used as negative control, based on the notion that mTOR activity is crucial in the developmental stages of the neurite network as well as for the synaptic strength at later stages [38]. For each compound, different concentrations were tested in unperturbed cultures to expose the dose range that does not exert negative effects in basal conditions. Concentrations were based on previously reported IC_50_ values from neuronal cell-based assays (Fig. 3a) [37,39-42]. To further reduce variability and aid legibility, scores of challenged conditions were normalized to their culture age-matched controls (Fig. 3b). Of all compounds tested, rapamycin had an overt negative impact on neuronal connectivity across the tested dose range. Conversely, the DLK inhibitor had an unequivocal positive effect at different culture ages (except for DIV 18), and this for doses below 1 µM. Higher concentrations induced neurotoxicity, which could be expected according to previously reported IC50 values [37]. These findings were also confirmed – albeit to a weaker extent - when using a trained RFC based on morphological descriptors to predict the connectivity degree (as culture time) of treated cultures (Additional file 8: Figure S7). Thus, the connectivity score revealed a pharmacological dose and time window in which DLK inhibition promoted connectivity in unperturbed cultures.

**Fig 3.**
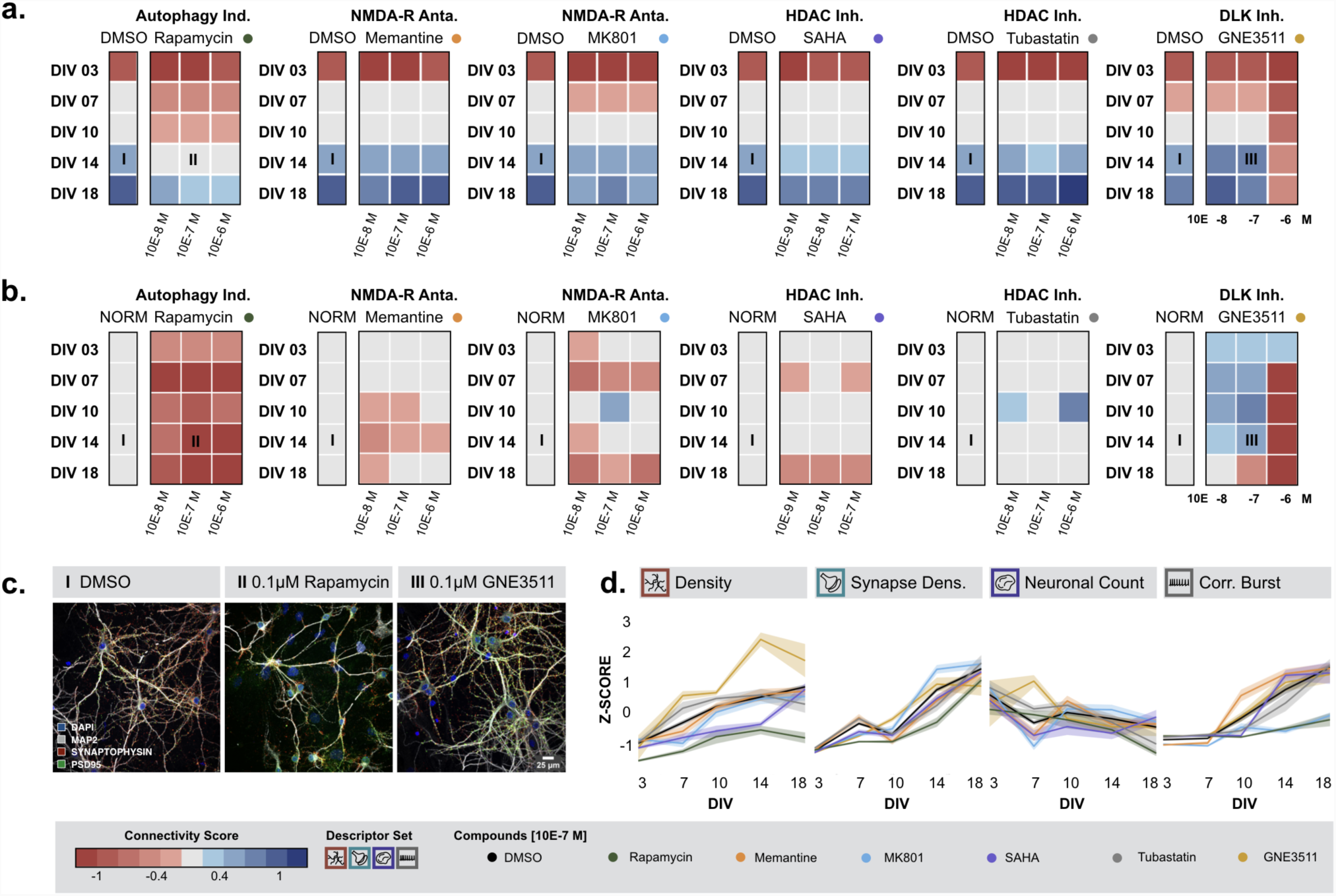
Focused compound screen identifies DLK as positive modulator of neuronal connectivity. **(a)** Connectivity scores of cultures continously treated with 3 different concentrations of selected compounds. **(b)** Connectivity scores based on z-scores normalized to their culture age-matched controls revealed that rapamycin is a strong negative modulator of neuronal connectivity and GNE3511 exerts a lasting positive effect on neuronal connectivity. (All compounds except GNE3511: Morph.: n_bio_=3 x n_tech_=5 – Func.: n_bio_=3 x n_tech_=6) (GNE3511: Morph.: n_bio_=2 x n_tech_=6 -- Func.: n_bio_=2 x n_tech_=9) **(c)** Example images of control cultures and cultures treated with 10E-7 M rapamycin or GNE3511 (14 DIV) (~I, II and III in panel b) (**d**) Z-scores showing the effect of the selected compounds on dendrite density, synapse density, neuronal count and the correlation of the calcium bursts. GNE4511 has a clear positive effect on dendrite and synapse density, whereas rapamycin and MK801 negatively affect the functional activity (mean and standard errors are visualized).

### DLK inhibition has both neuro-protective and -restorative potential

The observation that DLK inhibition exerted a positive effect on neuronal connectivity in primary cultures under basal conditions, drove us to test whether the same treatment was able to prevent or slow down the gradual connectivity loss observed in old cultures (Fig. 1, Fig. 4 and Additional file 9: Figure S8). To this end, we incubated cortical cultures with two different DLK inhibitors (GNE3511 and GNE8505) from DIV 21 onwards and followed them up to 68 DIV. A positive effect on the connectivity score was found in older cultures (>= DIV48) treated with a low dose (0.01 µM) of GNE3511 [37]. Yet, this effect was not consistently recapitulated with GNE8505 [39].

**Fig 4.**
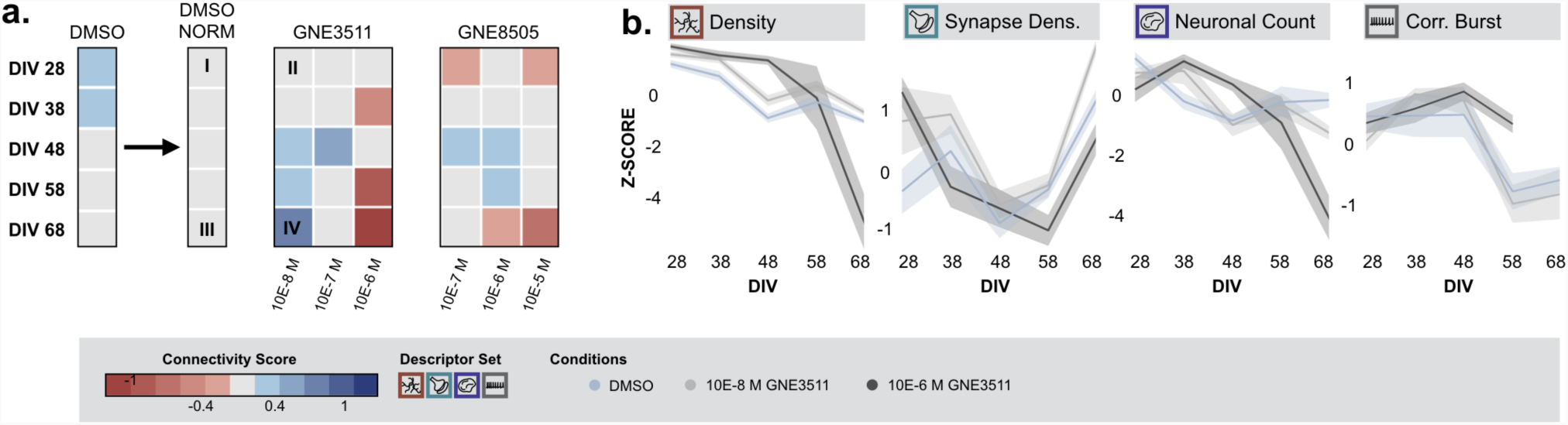
DLK inhibition can partially prevent the age related loss of connectivity. **(a)** GNE3511 (10E-8 M) could prevent the age-related loss of connectivity. Other effects were not lasting or neurotoxic (Morph.: n_bio_=1 x n_tech_=6 -- Func.: n_bio_=1 x n_tech_=9). **(b)** Z-scores of example descriptors. Treatment with 10E-8 M GNE3511 has a positive effect on the neurite density and synapse density. Treatment with 10E-6 M has however a negative impact on neuronal connectivity at later time points, indicated by a decreased neurite density and neuronal count (mean and standard errors are visualized)

Next, we tested whether DLK inhibition could also prevent cultures from degenerating when grown under sub-optimal or challenged conditions. Taking advantage of the fact that neurons do not dispose of well-established antioxidant protection mechanisms and are therefore highly susceptible to oxidative stress [40,41], we grew cortical cultures in medium without antioxidants (-AO). This had an adverse impact on neuronal connectivity, as reflected by a decreased connectivity score from 7 DIV onwards (Additional file 10: Figure S9). To limit the experimental load, we limited assessment to one time point, where the effect of antioxidant depletion was sufficiently clear, namely 14 DIV. Chronic treatment with 0.1µM GNE3511 could slightly improve the connectivity in comparison with the DMSO-treated cultures deprived from antioxidants (Fig. 5b), but the increase in the z-score of key morphological descriptors was not significant (p>0.05, pairwise Wilcoxon test with Bonferroni correction) (Fig. 5d). Chronic treatment with DLK inhibitor GNE8505 (0.1 µM and 1 µM) drastically improved the connectivity score in comparison with DMSO-treated-AO cultures (Fig. 5b,c). The z-scores of dendrite density, synapse density and neuronal count after treatment with 1µM GNE8505 showed no significant differences with those measured in unperturbed control cultures (p>0.05, pairwise Wilcoxon test with Bonferroni correction) (Fig. 5d). These results suggest a neuroprotective effect of GNE8505 in this compromised growth condition.

**Fig 5.**
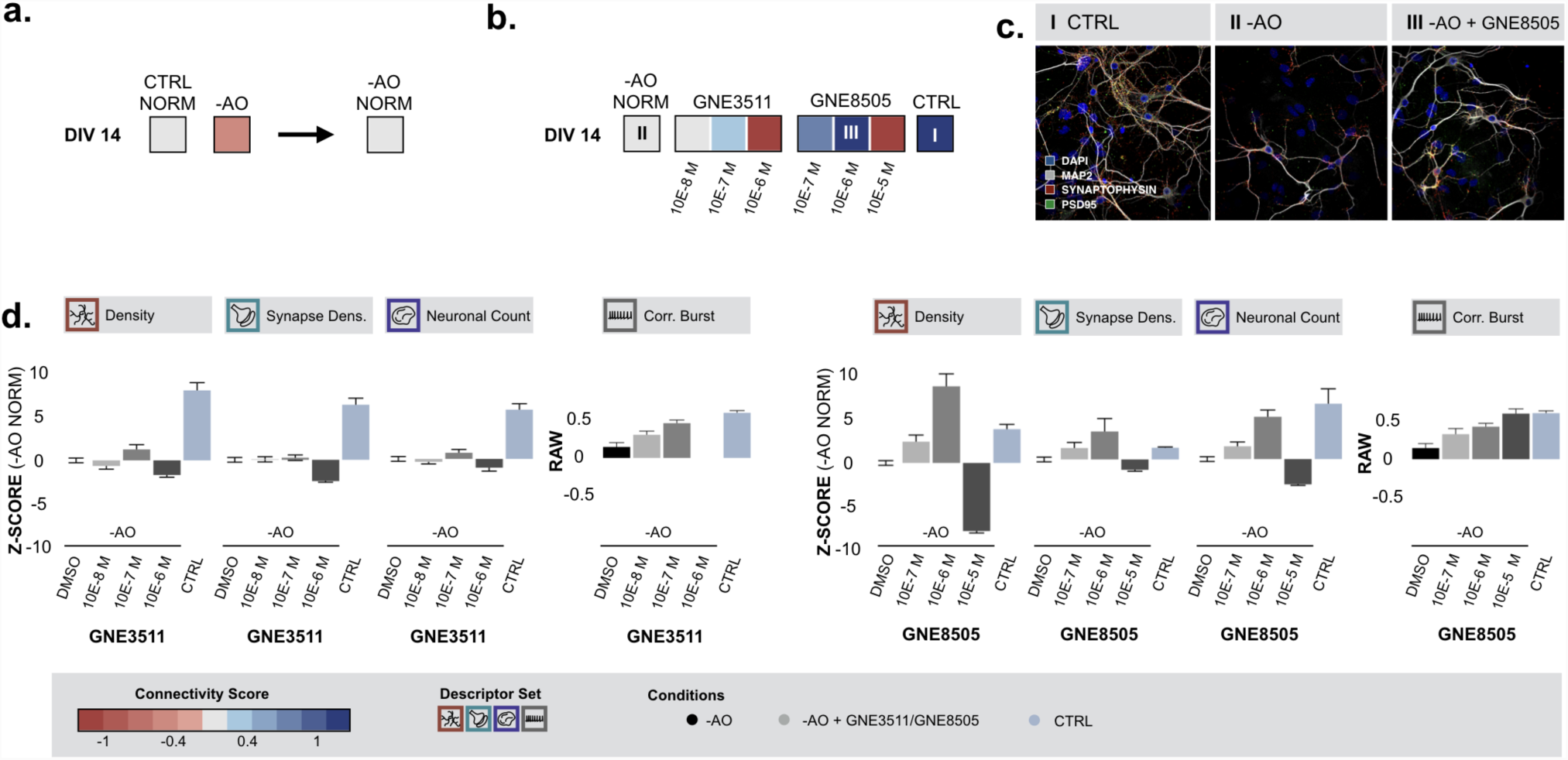
DLK inhibition has a neuroprotective potential in cultures deprived from antioxidants. **(a)** Primary cultures deprived from antioxidants (-AO), display impaired neuronal network connectivity at DIV 14 (Morph.: n_bio_=2 x n_tech_=6 -- Func.: n_bio_=2 x n_tech_=9). (**b**) Normalized connectivity scores of cultures deprived of antioxidants show that treatment from 0 DIV onwards with DLK inhibitors (GNE3511 and GNE8505) could prevent connectivity loss at 14 DIV (excl. highest concentration) (Morph.: n_bio_=2 x n_tech_=6 -- Func.: n_bio_=2 x n_tech_=9). (**c**) Representative images of control cultures, - AO cultures and -AO cultures treated with 10E-6 M GNE8505 (~I, II and III in panel b). (**d**) Z-scores normalized to -AO cultures of dendrite density, synapse density and the neuronal count. Treatment with 10E-6 M GNE8505 could retain the levels of control conditions. In addition to the morphological data, the raw data of the functional activity is plotted.

To assess the specificity of the protective action, we next introduced a completely different type of challenge to the cultures. We selectively altered microtubule stability through overexpression of hTau.P301L. Upon overexpression (at 3 DIV), a progressive decline in neuronal connectivity from 10 DIV onwards was witnessed (Additional file 10: Figure S9). When supplementing these challenged cultures with GNE8505 however, we found a strong positive impact on the connectivity score in comparison with DMSO treated cultures overexpressing hTau.P301L at DIV 14 (Fig. 6b). After treatment with 1µM or 10µM GNE8505, z-scores of dendrite density, synapse density (10µM only), and neuronal count no longer showed significant differences with untreated control cultures (p>0.05, Wilcoxon with Bonferroni correction). This indicated a broad-spectrum neuroprotective effect in challenged growth conditions (Fig 6c,d). The same trends were observed using the aforementioned morphological RFC approach (Additional file 11: Figure S10a). We also tested a pseudotime approach, typically used for single cell RNA sequencing results of cell differentiation [42-44], and found a distinct deviation from the stereotypical morphological trajectory (as built from control cultures) upon overexpression of hTau.P301L (Additional file 11: Figure S10b), which could be partially restored with GNE3511 treatment. Thus, using different algorithms based on different subsets of descriptors, we confirmed the neuroprotective effect of DLK inhibition in challenged conditions.

**Fig 6.**
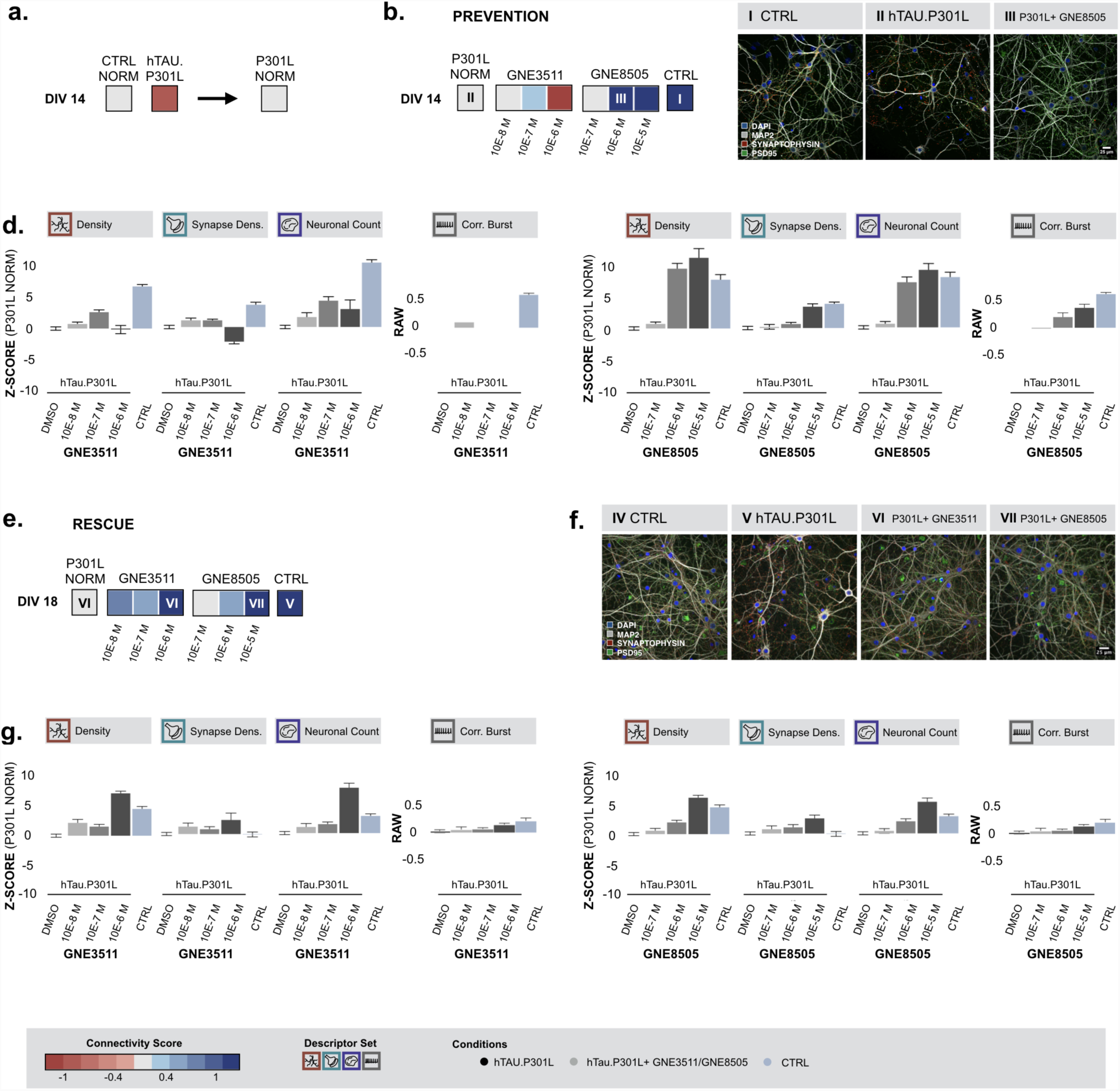
DLK inhibition has both neuro-protective and -restorative potential in cultures overexpressing hTau.P301L. (**a**) Cultures overexpressing hTau.P301L, showed impaired neuronal network connectivity (Morph.: n_bio_=2 x n_tech_=6 -- Func.: n_bio_=2 x n_tech_=9). **(b)** Continuous treatment with DLK inhibitors prevented loss of neuronal connectivity of the hTau.P301L model according to the normalized connectivity scores (Morph.: n_bio_=2 x n_tech_=6 -- Func.: n_bio_=2 x n_tech_=9). (**c**) Representative images of control cultures, -cultures overexpressing hTau.P301L and hTau.P301L cultures treated with 10E-6 M GNE8505 (~I, II and III in panel b). (**d**) Z-scores of morphological descriptors show that treatment with 10E-5 M GNE8505 could retain the levels of control conditions. Next to the morphological data, the functional correlation of the calcium bursts is plotted. **(e)** DLK inhibition rescues cultures overexpressing hTau.P301L after connectivity loss had already started (Morph.: n_bio_=1 x n_tech_=6 -- Func.: n_bio_=1 x n_tech_=6). (**f**) Representative images of control cultures, - cultures overexpressing hTau.P301L and hTau.P301L cultures treated with 10E-6 M GN3511 or 10E- 5 M GNE8505 (~IV, V, VI and VII in panel e). (**d**) Z-scores of morphological descriptors and the raw data of the correlation of the calcium bursts show a neuroprotective effect of both DLK inhibitors.

To test whether DLK inhibition could not just prevent but also reverse this process, we repeated the experiments in cultures overexpressing hTau.P301L, but this time we only started treating cultures with DLK inhibitors at a time point where connectivity loss was already manifesting (DIV14). Strikingly, in cortical cultures that overexpressed hTau.P301L, a rescue of connectivity loss could be observed at DIV 18 (Fig. 6e). After treatment of hTau.P301L-overexpressing cells with 10µM GNE8505, dendrite density was at the same level as for untreated control cultures (p>0.05, Wilcoxon with Bonferroni correction) (Fig. 6f,g). Treatment with 1µM GNE3511 even significantly increased the density area above control levels (p<0.05, Wilcoxon with Bonferroni correction) (Fig. 6f,g). Thus, we conclude that DLK inhibition is able to protect cortical cultures from diverse challenges and even partially rescue cultures from ensuing connectivity loss.

## Discussion

The continuous remodeling of the neuronal network is crucial for learning, memory and behavior, but is disrupted in several psychiatric and neurodegenerative disorders [4,45]. Identification of novel therapeutic targets requires a method that is able to quantify neuronal network connectivity over time with high accuracy and throughput. To our knowledge, the high-throughput analysis proposed in this paper is the first to comprehensively gauge neuronal connectivity in primary cultures by including descriptor sets reporting on the dendrite network, synapse markers, nuclei and calcium bursting activity. Other high-throughput studies aimed at finding regulators of neuronal connectivity [17-21], have mainly focused at one or few readouts such as neuron number [19] and neurite outgrowth [18,19], synapse density [22,23] or calcium responses [29]. The strength of an integrative approach lies in the fact that it can account for a number of potential sources of bias. For instance, when only considering synapse density, observed changes could be the result of true variations in the synapse number with equal neurite density [20,21], but they could equally arise from an altered neurite density with preserved synapse count [26], an increased density of only one or both synaptic partners (pre/post) [20], an increased spot size of one or both synaptic partners or an altered clustering of neuronal somata and fasciculation of neurite bundles. Furthermore, the inclusion of calcium data as a proxy for spontaneous electrical network activity allows determining whether alterations in the network morphology also have a functional impact and vice versa. For example, previously published data by our group shows that overexpression of human tau in hippocampal cultures decreased the neurite density but increased the synaptophysin density [26]. At a functional level, the percentage of active neurons and frequency of synchronous bursts decreased while the synchronicity was preserved. This underscores the argument that conclusions may vary depending on the descriptors used and that an integrative approach is desirable.

In line with other studies, we found an increase in dendrite density and synapse density with culture age, and - although not linear – a strong increase in the synchronous bursting behavior of the network at later time points (>10 DIV). The time scales used in literature range from DIV6 to DIV21, which corresponds with the time range we identified in our initial experiment to be the time span in which dendrite density, synapse density and functional activity increased drastically. The decrease in neuronal cell count with culture age also confirmed previous findings in which many neurons die during culture, but the surviving neurons form a network [21,46]. To our knowledge no other studies have measured descriptors of both neuronal and glial nuclei in the context of a connectivity study. We showed however that certain treatments (e.g. AraC) had a major effect on nuclear descriptors.

When applying PCA to the unfiltered morphological descriptor set, we unveiled distinct temporal trajectories [47], suggesting that the culture age might serve as an indicator of connectivity. This was confirmed by the fact that a random forest classifier could predict culture age with an accuracy of 95%. Yet, to also include functional parameters we defined a connectivity score. Using this score as readout, we could confirm that inhibition of mTOR activity, using rapamycin, impairs the development of the neurite network [38,48]. Nevertheless, rapamycin has been used to slow down or block neurodegeneration in mouse models of Alzheimer’s, Parkinson’s and Huntington’s disease, through the induction of autophagy that cleared accumulated autophagosomes and/or aggregated proteins in these models [49-51]. Therefore, an extension of this work could focus on assessing the effect of rapamycin on neuronal connectivity under conditions of induced toxic protein aggregation. Antagonists of NMDA-R, such as memantine and MK801, were also tested. These antagonists have the potential to block the excessive NMDA-R activity in many neurodegenerative diseases, hereby reducing the increased calcium influx in the neurons [32,33]. NMDA-R antagonists can however also block the normal function of these receptors, which was suggested by the overall negative effect on the connectivity score for MK801 (in which the functional descriptors had a major contribution). This score also indicated that memantine treatment had only a slight negative impact on neuronal connectivity in comparison with MK801, which may be the result of memantine having a shorter dwell time on the NMDA-R then MK801 [32]. Tubastatin, a HDAC inhibitor that improved cognitive deficits in mouse models of Alzheimer Disease by improving microtubule stability [34], did not have a negative, nor a neurotrophic, effect on the overall neuronal connectivity in control conditions, and may therefore be further explored in compromised conditions. Treatment with another HDAC inhibitor, SAHA, resulted in a decreased neuronal connectivity at later time points. This may be due to the fact that SAHA targets multiple HDAC classes, whereas tubastatin only targets HDAC6 [52].

The sole compound that showed a clear neurotrophic effect in unperturbed primary cultures was an inhibitor of DLK, which is why we chose to pursue the efficiency of this compound in compromised conditions. DLK, also known as MAP3K12, is an upstream regulator of the c-Jun N-terminal kinase (JNK) stress response pathway, which becomes activated in both acute and chronic neurodegenerative conditions_[53]_. Its activation induces a broad transcriptional injury response (via c-Jun and ATF4) [54]. The fact that it is upstream in this highly conserved pathway makes that DLK inhibition has a broad action range, at least in preclinical models. Pharmacological or genetic DLK inhibition protects against excitotoxicity [55], growth factor deprivation [56-58], Amyloid and Tau pathology [36,59], nerve crush and traumatic brain injury [54,60] retinal ganglion cell degeneration [57,61] and SOD1-mediated neurodegeneration [36]. At this moment, one inhibitor is in Phase I clinical trial for Amyotrophic Lateral Sclerosis (ALS) by Genentech (Roche) [62]. Our results now show that also in unperturbed primary cortical cultures DLK inhibition results in enhanced morphofunctional connectivity. Yet, it should be noted that the inhibitor treatment started 4 hours after plating. It is possible that the cells already experienced stress of the dissociation procedure, which could have been attenuated by DLK inhibition.

We found that DLK inhibition could prevent and even rescue neurodegeneration induced by hTau.P301L overexpression. This aligns with literature data where double mutant mice (Tau^P301L^;DLK^cKO^) show attenuated cell loss the subiculum compared to single mutant (Tau^P301L^) mice without having altered tau pathology [36]. Our *in vitro* approach now also shows that hTau-P301L-induced neuron loss is accompanied by impaired neurite formation, synapse density and functional activity and that this can be prevented and rescued by pharmacological DLK inhibition. According to the connectivity score, the neuroprotective effect of DLK inhibitor GNE8505 was higher compared to GNE3511. This could be explained by the higher kinase selectivity of GNE8505 [39].

The omission of anti-oxidants from the culture medium of primary cortical neurons resulted in gradual impairment of morphofunctional connectivity, which could be prevented by DLK inhibition as well. Neurons are particularly sensitive to reactive oxygen species (ROS) and most neurodegenerative disorders are associated with increased oxidative stress [40], making our *in vitro* model attractive for drug screening. In line with our results, it was described before that the SOD1^G93A^ mouse model for ALS exhibits neuronal cell death due to an impaired oxidative stress defense and that this is accompanied by aberrant JNK pathway activation [36]. Double mutant mice (SOD1^G93A^;DLK^cKO^) show enhanced neuronal survival and myelinization as well as reduced neuroinflammation compared to single mutants (SOD1^G93A^), resulting in enhanced grip strength and longer life span of the former model.

Finally, we also verified if DLK inhibition could sustain morphofunctional connectivity in aging cultures, as it is known that JNK signaling is elevated in the aging brain [63,64]. However, the effect was rather limited. Increased variability across aged cultures and reduced sensitivity of the morphological readout due to the very high density of neurites covering most of the culture plate could have masked a potential clearer effect of DLK inhibition on ageing. To increase the sensitivity of the current approach, it could be advantageous to include more synapse markers so as to better map the full landscape of synapse types (e.g. inhibitory vs excitatory). To bypass spectral limitations, one could resort to the use of narrow-emission band labels, such as Quantum Dots (QD) [65], or divert to cyclic immunofluorescence staining protocols [66],[67]. The latter would however lower the throughput and put a higher demand on downstream analyses (*e.g.*, image registration). A particular caveat of the current approach was the high variability in functional readout. To strengthen the sensitivity of this assay one could resort to imaging more and larger cell populations (field of view). Previous studies have revealed the co-existence of separate networks of high connectivity within the large neuronal network that may not always fire in sync [68,69] Identification of stratified connectivity patterns may therefore expose more subtle modifications of the functional connectivity. Selective addition of chemical stimuli (e.g. glutamate) could further unveil cell type specificity as well as differences in spontaneous and induced functional activity [30].

## Conclusion

In conclusion, we have shown that morphofunctional profiling of primary cultures using deep coverage microscopy allows accurate quantification of neuronal connectivity *in vitro*. We established a connectivity score, including morphological and functional correlates that was used to identify modulators of neuronal connectivity. With our approach we were able to expose a dose and time window for DLK inhibitors that evoked positive effects on neuronal connectivity and could even rescue perturbed cultures. Therefore, the current approach holds promise for identifying pathways and treatments that preserve or rescue neuronal connectivity in neurodegenerative disorders.

## Methods

### Preparation of primary neuronal cultures and pharmacological treatment

This study was carried out in accordance with the recommendations of the ethical committee for animal experimentation of the University of Antwerp (approved ethical file 2015-54). Hippocampi and cortex were dissected from WT E18 C57Bl6 mouse embryos in Hepes (7 mM)-buffered Hanks Balanced Salt Solution, followed by trypsin digestion (0.05%; 10 min; 37°C) and mechanical dissociation by trituration through 2 pipette tips with decreasing diameter. After centrifugation (5 min at 200g), the cell pellet was resuspended in Minimal Essential Medium supplemented with 10% heat-inactivated normal horse serum and 30 mM glucose. Cells were plated in Poly-D-Lysin-coated 96-well plates (Greiner Cell coat, µClear), at 30,000 cells/cm^2^, and kept in a humidified CO_2_ incubator (37°C; 5% CO_2_). After 4 hours, the medium was replaced with B27 (2%) supplemented Neurobasal medium, containing Sodium Pyruvate (1 mM), Glutamax (2 mM), glucose (30 mM) and PenStrep (0.5%). For antioxidant deprivation, the commercially available B27 supplement minus antioxidants was used. To suppress proliferation of non-neuronal cells, 1 µM arabinosylcytosine was added in 25 µl Neurobasal-B27 medium at the third day after plating. Cell culture supplies were purchased from ThermoFisher. The following compounds were obtained from the in house Janssen pharmacy unless stated otherwise: rapamycin (Santa Cruz sc-3504), memantine, MK801, SAHA, tubastatin, GNE-8505, GNE-3511. These compounds were added in 25 µl Neurobasal-B27 medium at final concentrations of 0.01/0.1/1/10 µM. Control cultures received the same volume DMSO, which was always below 0.1% (Fig. 7).

**Fig 7.**
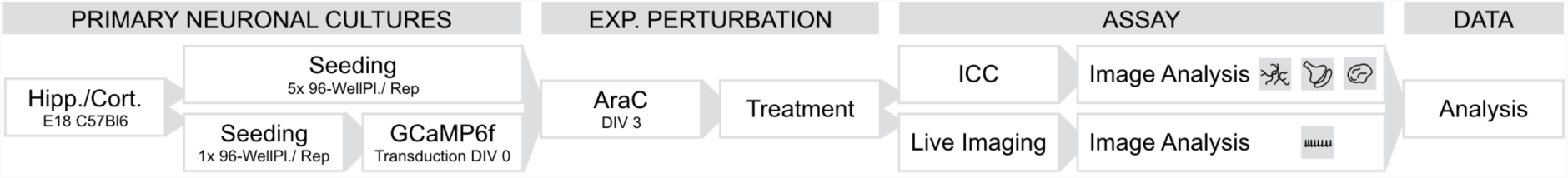
General workflow. General workflow of the microscopy-based framework. Hippocampi and/or cortex from WT E18 C57Bl6 mouse embryos are dissected, after which cell suspensions were created and seeded in 96 well plates. Cultures that would be used in the functional assay were transfected with an AAVDJ-hSyn1-GCaMP6f-nls-dTomato vector on DIV 0. At 3 DIV, cultures were treated with arabinosylcytosine (AraC) to suppress the excessive growth of non-neuronal cells. Cultures were treated every three or four days. At fixed time points, the morphological and functional characteristics of the cultures were quantified on which data analysis was performed.

### Immunocytochemistry

Cultures were fixed (2% PFA, 20 min, RT) at 3/7/10/14/18 DIV (or later DIV in experiments with an extended time range) and immunocytochemically labeled for a neurite marker (Chicken Polyclonal against MAP2, Synaptic Systems 188006, 0.5 µg/ml), a presynaptic marker (Guinea Pig Polyclonal against Synaptophysin-1, synaptic systems 101004, 0.5 µg/ml), a postsynaptic marker (Mouse Monoclonal against PSD-95, ThermoFisher MA1-046, 2 µg/ml) and a nuclear marker (DAPI, 5 µg/ml) (Fig. 7). In brief, the cultures were permeabilized with 1% Triton X-100 in blocking buffer (0.1% bovine serum albumin and 10% normal horse serum in PBS) for 10 minutes, followed by an overnight incubation with the primary antibodies at 4°C in blocking buffer. After washing with PBS, secondary antibodies (Donkey-anti-Chicken-Cy5 / Donkey-anti-GuineaPig-Cy3 / Goat-anti-Mouse-AlexaFluorPlus488, 1 µg/ml) were added for 2 hours at room temperature. Finally, DAPI was applied to the cultures for 10 minutes at a concentration of 2.5 µg/ml, followed by a PBS wash.

### AAV-mediated expression of GCaMP6f and MAPT-P301L

Functional connectivity was assessed by means of live cell calcium imaging of spontaneous neuronal activity (Fig. 7). At 0 DIV, a genetically encoded calcium indicator (GCaMP6f [27,29]) along with a nuclear-localized red fluorescent protein (nls-dTomato) was introduced via AAV-mediated expression under the synapsin promoter (addgene plasmid #51085 deposited by Jonathan Ting was packaged in-house with the AAV-DJ Helper Free Packaging System of Cell Biolabs; 0.01 µl crude lysate/well). Overexpression of *hTau.P301L* was induced via AAV-mediated transduction of an expression vector under the synapsin promotor [70,71] and were administered at 3 DIV at MOI 300.

### Image acquisition

Confocal images of immunostained cultures were acquired on an Opera Phenix High Content Screening System (40XW, NA 1.2, PerkinElmer). Per well, 15 fields were acquired in 4 channels (405 nm, 488 nm, 561 nm and 640 nm excitation) in 5 axial positions separated by a 1 µm spacing. Different fluorescence channels were separated using standard excitation/emission filters and dichroic mirrors. Calcium imaging was performed on a spinning disk confocal microscope (20X, NA 0.75, UltraVIEW VoX, PerkinElmer) at 37°C and 5% CO_2_. For each field, a 3 minute recording (2 frames per second) of the calcium activity was acquired in the 488 nm channel, followed by a Z-stack of the nuclear nls-dTomato signal in the 561nm channel. Per well, 3 fields were imaged.

### Image processing and analysis

Image analysis was carried out in Acapella® (PerkinElmer), but a similar pipeline is available for FIJI [72,73]. The acquired images were read in per field of view. After maximum projection of the z-stacks obtained from the MAP2 and DAPI channel, the nuclei were detected using a manually assigned threshold. Dendrites were identified using a rough (user-defined threshold) and fine (user-defined threshold after Frangi filtering [74]) segmentation. Neuronal nuclei were distinguished from non-neuronal based on a user-defined maximal projected area, minimal circularity and minimal occupancy in the dendrite mask. For both the dendrite network and nuclei a broad range of morphological and textural (object- and image-based) descriptors were extracted (Additional file 1: Table S1).

Next, both the dendrite mask and the neuronal nuclei mask were dilated and subtracted from each other to obtain a search region (i.e. dilated dendrites without neuronal nuclei) in which the pre- and postsynaptic spots were detected. The sharpest slice (based on the highest standard deviation of the intensity) from the presynaptic channel and its matched postsynaptic slice were used for spots detection. The spots were first enhanced using a difference of Gaussian filter with a user-defined kernel size, after which a user-defined threshold was applied to segment the spots. Pre- and postsynaptic spots that had an overlap of minimum 1 pixel were identified as a synapse. In addition to this object-based colocalization, the Pearson correlation of the pre-and postsynaptic channel was calculated as an intensity-based colocalization metric. Next to morphological and textural descriptors, the density of pre- and postsynaptic spots and synapses were calculated (Additional file 1: Table S1).

Calcium recordings were analyzed using a home-written MATLAB script, adapted from [30]. Briefly, the neurons were segmented based on the nls-dTomato nucleus image, after which traces of the GCaMP6f fluorescence intensity over time were generated. After normalization, signal analysis returned parameters such as percentage of active neurons, frequency and amplitude of (synchronous) calcium bursts and burst correlation, which is the average of the Pearson’s correlation matrix between all neuron pairs in the field of view.

### Data integration and representation

Data analysis and representation was done in R [75]. CSV files containing the field data were read and merged with the metadata according to the plate layout. Fields from the morphological assay lacking nuclei or dendrites were removed. False nuclear segmentations were identified based on projected nuclear areas that exceeded the mean projected area with 5 standard deviations. After filtering of the data, well averages were calculated for the morphological data and were depicted as technical replicates. Data extracted from the functional assay was not averaged to not further reduce the number of data points and each recording was seen as technical replicates as such. Z-scores of all descriptors were calculated using the mean and standard deviation within each experiment and replicate. PCA analyses were done on these z-scores using the *prcomp* function in R. Classification was done on the raw data using the *randomForest* function in R. Both the control cultures of morphological and functional data were split in a training (2/3) and test (1/3) set. For each classifier the optimal number of trees and descriptors was determined using 10-fold cross validation, after which the optimal settings were used to classify the test set.

The calculation of the connectivity score consisted of four steps; 1) first, the correlation of all descriptors with the culture age was determined for all control conditions and averaged across six separate experiments; 2) next, the inter-correlation of the descriptors was calculated in a merged dataset including control conditions and treated cultures; 3) descriptors were then ranked according to their average correlation with culture age. This ordered descriptor list, was then filtered top-down, by removing those descriptors with an absolute inter-correlation higher than 0.75 (Additional file 6: Table S5). Channel descriptors that were susceptible to outliers (e.g. intensity metrics) were removed as well; The final connectivity score was calculated as the average z-score of the subset of descriptors using their respective correlation with the culture age as weights. When comparing control cultures with perturbed or treated cultures, normalized z-scores were used based on the to the mean and standard deviation of control or untreated suboptimal conditions within each experiment, replicate and time point.

To construct the morphological trajectories, analogous to the cell differentiation trajectories based on single-cell RNA sequencing data, the R packages *scater* [76], *monocle* [42-44] and *singleCellExperiment* were used. All descriptors of control cultures were rescaled between 0 and 1, within each separate experiment. On the rescaled data, we applied the *differentialGeneTest monocle* function fitted to the DIV to order the descriptors by most to least significant. Finally, the standard deviation at DIV 18 for the control was calculated. Descriptors that were consistent during culture maturation should give the least dispersion at DIV 18. Based on those two measures, 37 descriptors were selected (q-value < 10E-20, standard deviation <1) that were used to build the morphological trajectories.

All the data representations were built using the *ggplot* function in R.

## List of abbreviations

AraC: arabinosylcytosine
DIV: days in vitro
DLK: dual leucine zipper kinase
HDAC: histon deacetylase
MCR: misclassification rate
NMDA-R: N-methyl-d-aspartate sensitive glutamate receptor
PCA: principal component analysis
RFC: random forest classifier
SAHA: suberoylanilide hydroxamic acid

## Declarations

### Ethics approval and consent to participate

The preparation of primary neuronal cultures was carried out in accordance with the recommendations of the ethical committee for animal experimentation of the University of Antwerp (approved ethical file 2015-54).

### Consent for publication

Not applicable.

## Authors’ contributions

PV, MV, RN, PL, AE, MV* and WDV conceived and designed the experiments. PV, IC, GG and EC performed the wet lab work. MV and WDV devised the image analysis script for neuronal morphology. PV analyzed the imaging data (morphological and functional) with available scripts. MV, PV, GG and WDV performed data analysis. PV, MV and WDV drafted the manuscript. All authors critically revised the manuscript and are accountable for all aspects of the work. (MV: Marlies Verschuuren, MV*: Mieke Verslegers)

## Acknowledgements

The authors would like to thank Sofie Thys for technical assistance during preparation of primary neuronal cultures.

## Additional files

Additional file 1: **Table S1.** All used descriptors of the dendrite, synapse, nuclear and functional descriptor sets. Measurements are averaged for every field of view.

Additional file 2: **Figure S1.** Differences between hippocampal and cortical cultures. Dendrite width and postsynaptic intensity are higher in hippocampal cultures. Nuclei of cortical cultures show significant textural differences with nuclei from hippocampal cultures. The neuronal ratio of cortical nuclei is higher than the ratio in hippocampal cultures, leading to an overall decrease in the projected area of the nuclei.

Additional file 3: **Figure S2**. Cultures can be clustered and accurately classified. **(a)** Representative images of both hippocampal and cortical cultures at different DIV for 3 different replicates. **(b)** PCA based on different descriptor sets. Nuclear descriptors entail information about the culture type based on the ratio of non-neuronal versus neuronal nuclei and textural descriptors of the nuclei. PCA of synaptic descriptors discriminated between different time points. The clustered time points formed phenotypic trajectories that progress with increasing synapse density. PCA based on dendrite descriptors resulted in the best clustering results based on only one descriptor class, but the difference between replicates was also most pronounced in this representation. The integrated morphological assay outperformed the functional assay in clustering both culture age and types (n_bio_=3 x n_tech_ =6). **(C)** Confusion matrices of classification results using a random forest classifier. The misclassification rate (MCR) was lowest using the integrated morphological descriptor set (n_bio_=3 x n_tech_ =6).

Additional file 4: **Figure S3**. Functional descriptors entail unique information about neuronal connectivity. MK801 treatment (yellow) impaired the functional activity at 14 and 18 DIV, that was not reflected in the morphological data (Morph.: n_bio_=3 x n_tech_=5 – Func.: n_bio_=3 x n_tech_=6) (mean z-scores and standard errors are visualized).

Additional file 5: **Figure S4**. Nuclear descriptors entail unique information about neuronal connectivity. AraC treatment (yellow) had a major negative impact on nuclear descriptors during the whole time range, while other descriptors showed only transient effects (e.g. dendrite density) or negative effects on later time points (e.g correlation of the calcium cursts) (Morph.: n_bio_=3 x n_tech_=6 – Func.: n_bio_=3 x n_tech_=6) (mean z-scores and standard errors are visualized).

Additional file 6: **Figure S5**. Descriptor selection. A subset of descriptors was selected based on the calculated correlation with the DIV and inter-correlation between each descriptor. First the descriptors were ranked according to their correlation with the culture age. This rank was used to subsequently add (blue) descriptors to the final subset, making sure that the inter-correlation within descriptors of the subset did not exceed 0.75. Image based descriptors were excluded from the descriptor set, because they were sensitive to outliers.

Additional file 7: **Figure S6**. Connectivity scores are sensitive to changes in denrite, synapse, nuclear and functional descriptors. **(a)** Connectivity scores of MK801 treated cultures showed greater differences with DMSO treated cultures at later time points when based on the integrated dataset in comparison with the ones based on only morphological data. (Morph.: n_bio_=3 x n_tech_=5 – Func.: n_bio_=3 x n_tech_=6). **(b)** Connectivity scores of AraC treated cultures revealed greater connectivity impairments in comparison with DMSO treated cultures when including nuclear descpriptors (Morph.: n_bio_=3 x n_tech_=6 – Func.: n_bio_=3 x n_tech_=6).

Additional file 8: **Figure S7**. Classification of morphological data confirms findings based on connectivity score. A random forest classifier that was trained on pooled DMSO treated cultures revealed a negative impact of rapamycin on neuronal connectivity as can be seen from the cultures that were misclassified and were assigned a culture age that was less than the actual culture age (red). Treatment with 0.01µM and 0.1 µM of GNE3511 could however improve the neuronal connectivity (~green) (Morph.: n_bio_=2 x n_tech_=6).

Additional file 9: **Figure S8**. Extended culture age reveals slight reduction of neuronal connectivity. Connectivity scores based on z-scores from cortical cultures grown for an extended period of time. Neuronal connectivity increased drastically during the first two weeks, after which it stagnated up to five and a half weeks. From DIV 38 onwards age-related loss of connectivity was detectable (Morph.: n_bio_=1, n_tech_=6 -- Func.: n_bio_=1, n_tech_=9).

Additional file 10: **Figure S9.** Impaired neuronal connectivity in compromised models. **(a)** Primary cultures deprived from antioxidants (-AO), had a negative impact on neuronal network connectivity form 7 DIV onwards (Morph.: n_bio_=2 x n_tech_=6 -- Func.: n_bio_=2 x n_tech_=9). **(b)** Cultures overexpressing hTau.P301L, showed a decreasing neuronal connectivity from 10 DIV (Morph.: n_bio_=2 x n_tech_=6 -- Func.: n_bio_=2 x n_tech_=9).

Additional file 11: **Figure S10.** Alternative data analyses confirm findings based on connectivity score. **(a)** A random forest classifier that was trained on pooled DMSO treated cultures confirmed the negative impact of hTau.P301L overexpression on later timepoints (red). Chronic treatment of those cultures with 0.01µM and 0.1 µM GNE3511 could however improve the neuronal connectivity in the suboptimal conditions (shift from red to green). (**b**) Morphological trajectories build based on pooled DMSO treated cultures revealed. hTau.P301L overexpression induced a shift of these cultures into a side branch of the trajectory. Treatment with 0.01µM or 0.1 µM GNE3511 could prevent this shift (Morph.: n_bio_=2 x n_tech_=6).

