## Supplementary figures and images for "Deep coverage microscopy exposes a pharmacological window for modifiers of neuronal network connectivity"

### Additional file 2: Figure S1

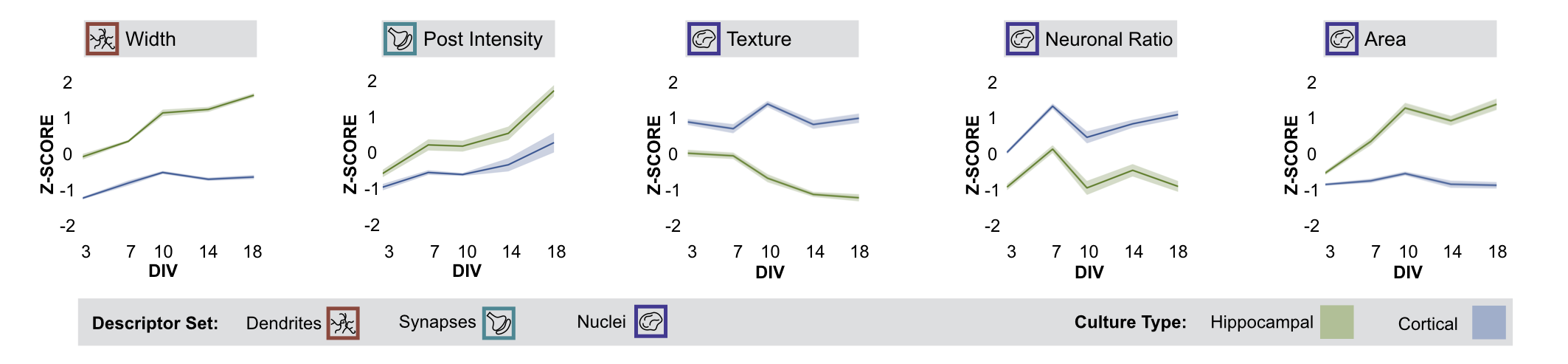

### Additional file 3: Figure S2

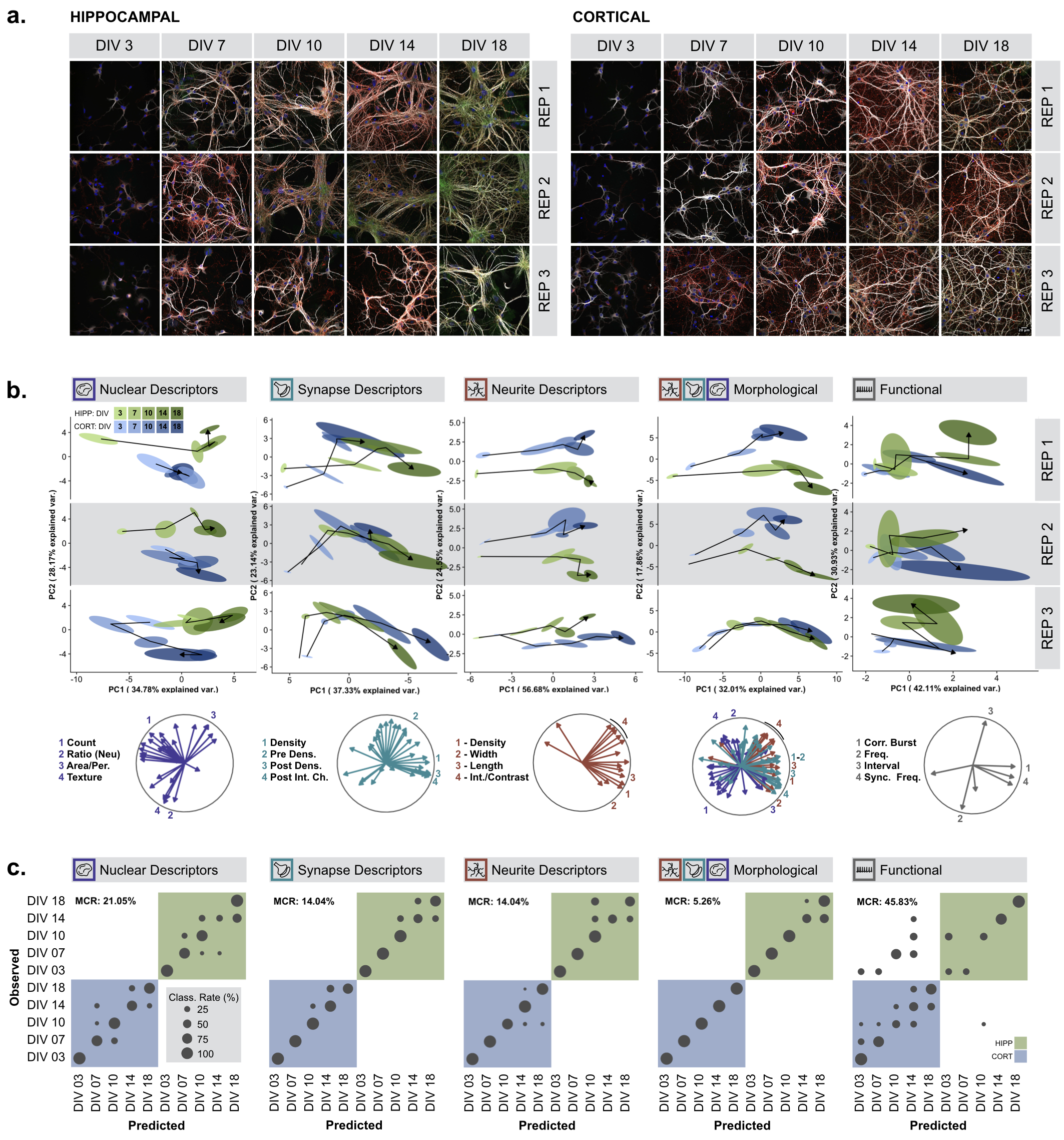

### Additional file 4: Figure S3

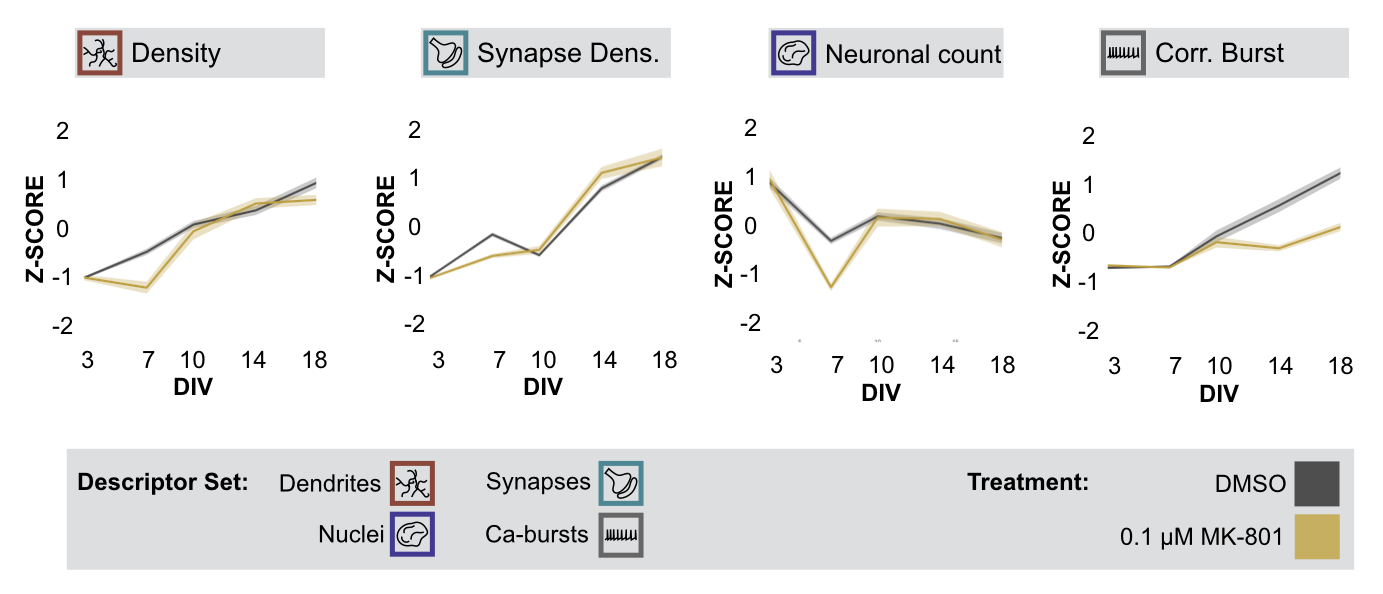

### Additional file 5: Figure S4

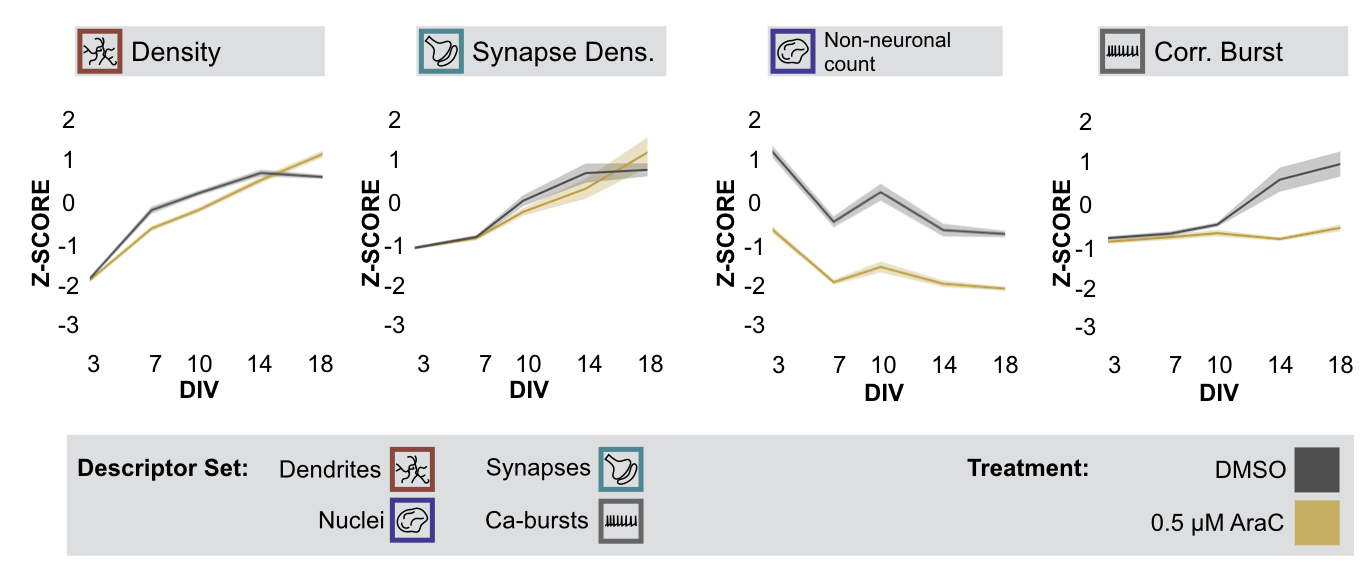

### Additional file 6: Figure S5

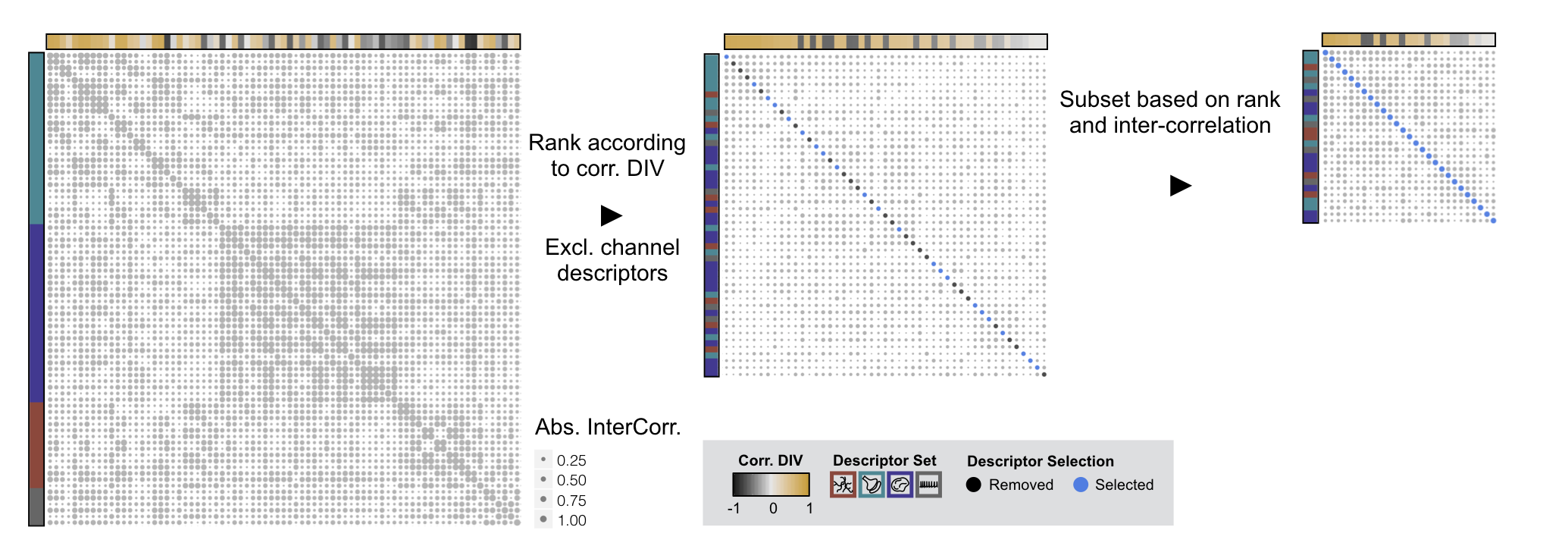

### Additional file 7: Figure S6

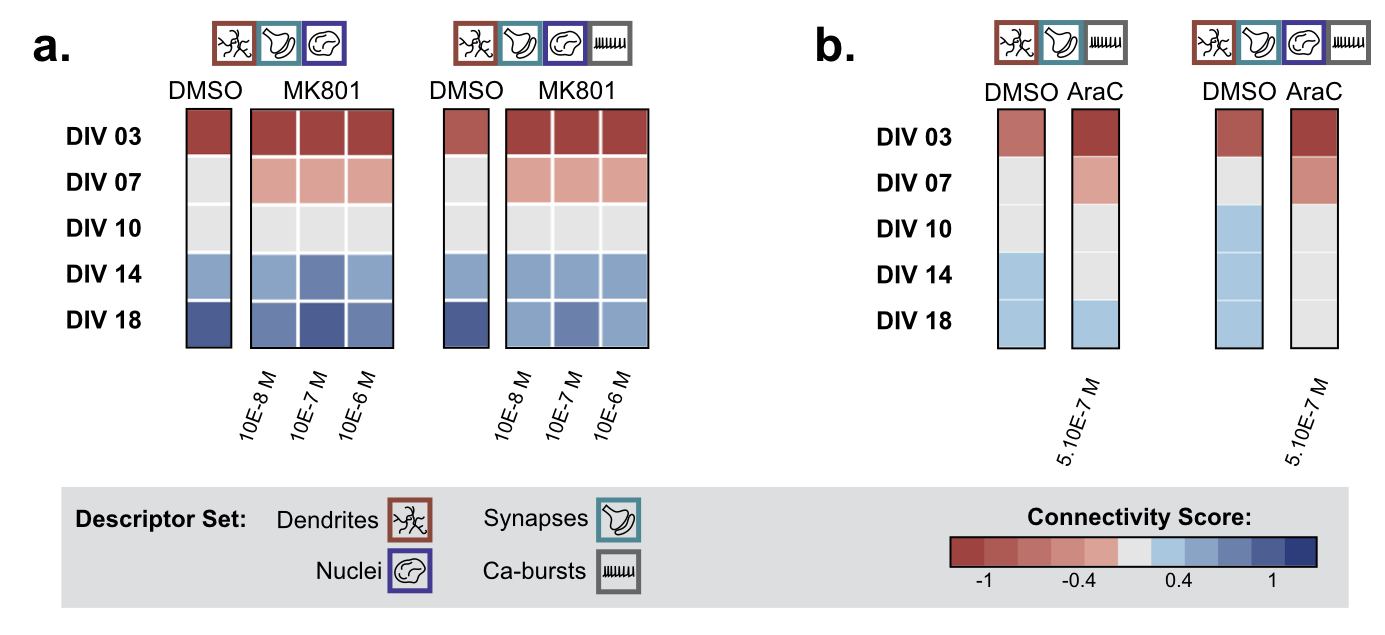

### Additional file 8: Figure S7

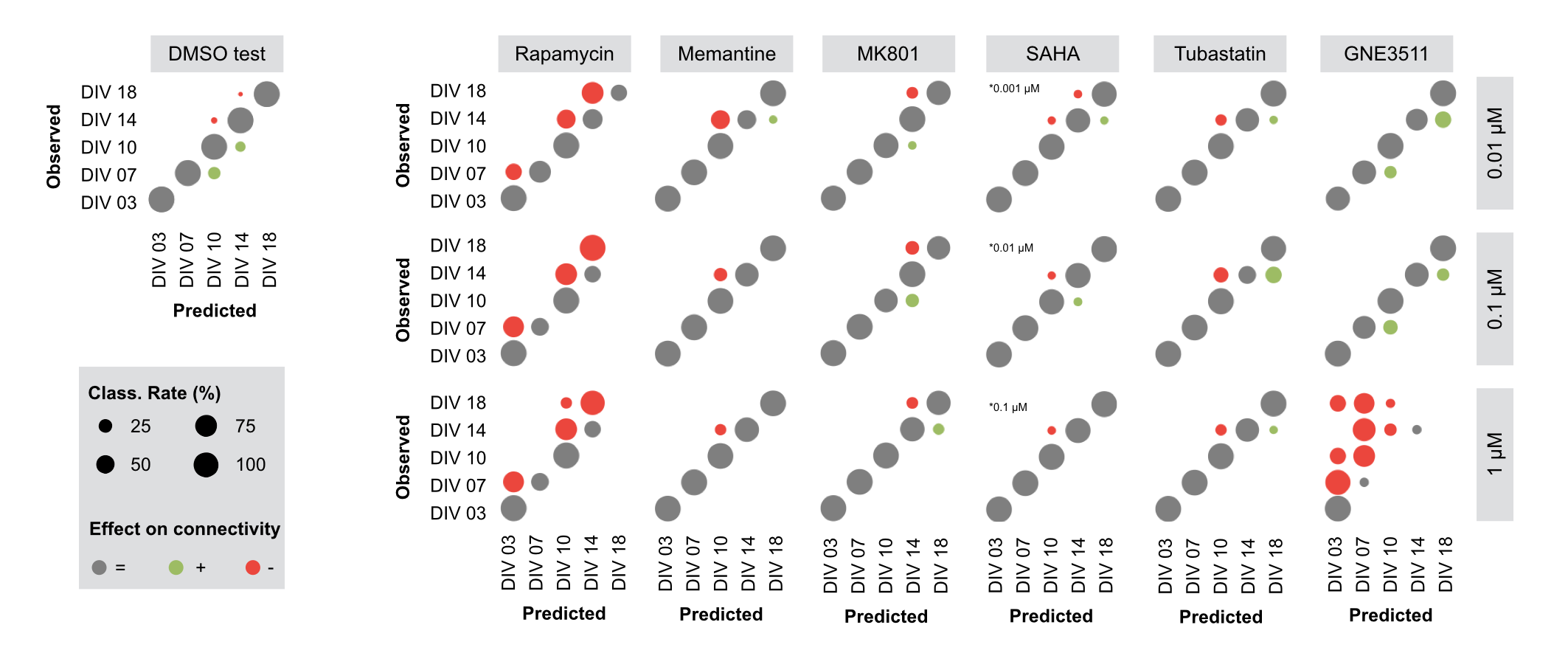

### Additional file 9: Figure S8

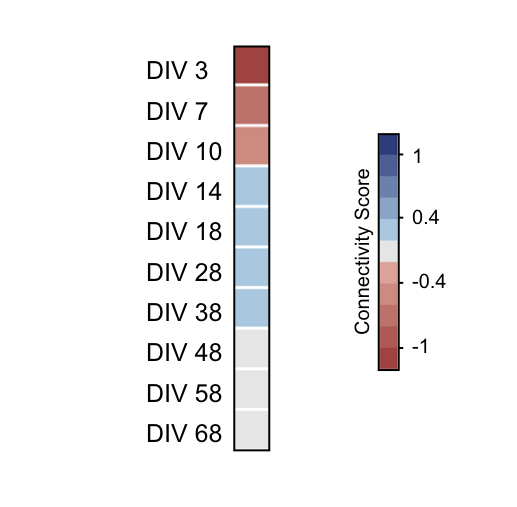

### Additional file 10: Figure S9

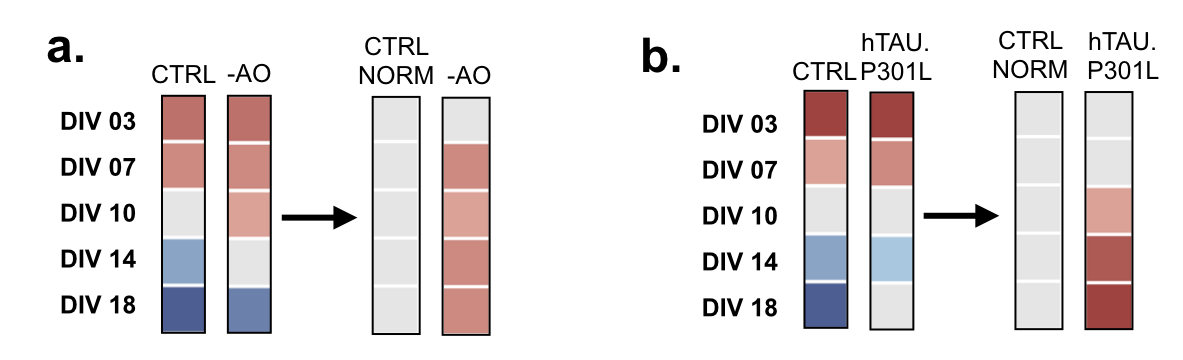

### Additional file 11: Figure S10

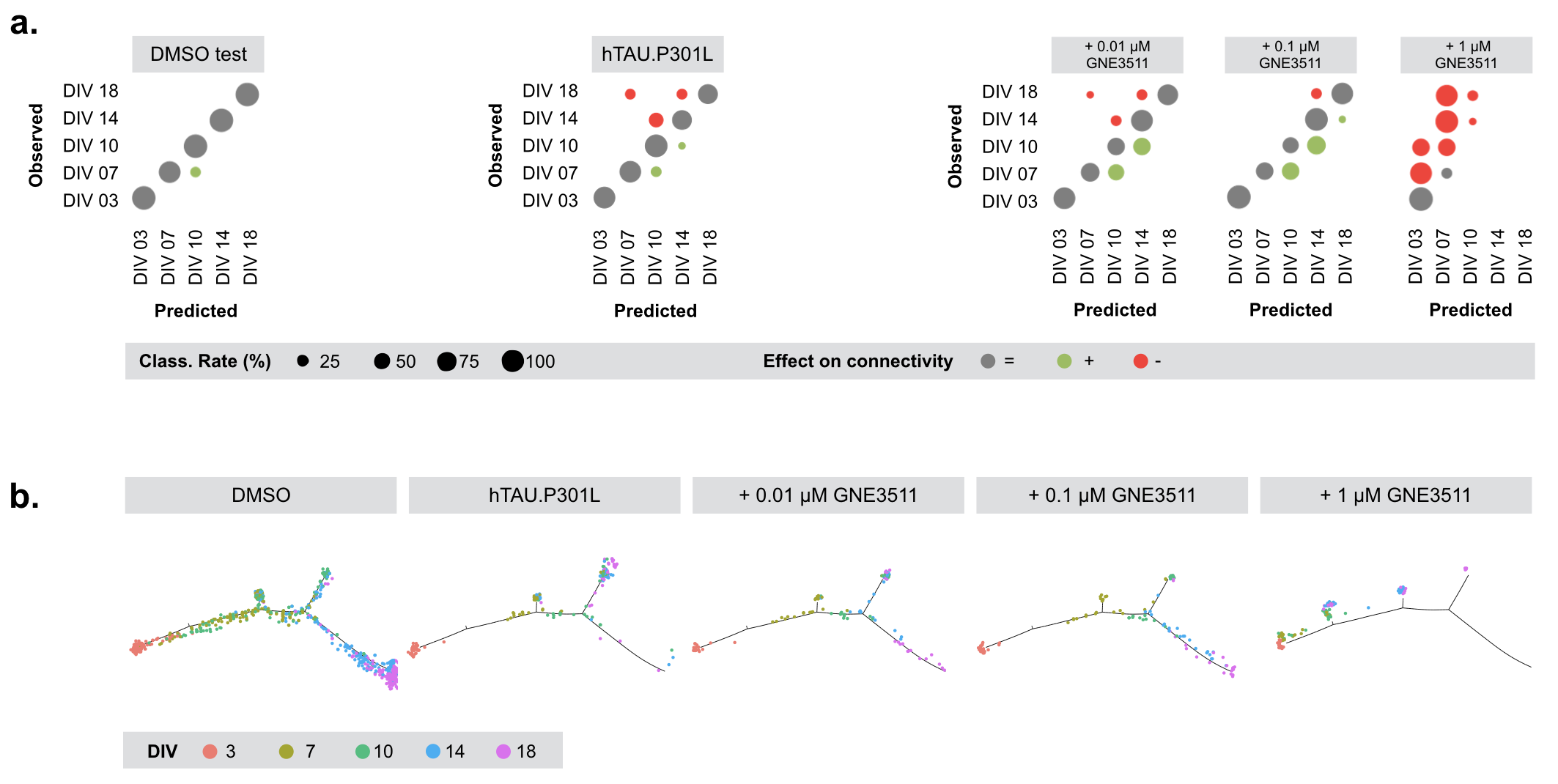
